# dynDeepDRIM: a dynamic deep learning model to infer direct regulatory interactions using single cell time-course gene expression data

**DOI:** 10.1101/2021.08.28.458048

**Authors:** Yu Xu, Jiaxing Chen, Aiping Lyu, William K Cheung, Lu Zhang

**Affiliations:** Department of Computer Science, Hong Kong Baptist University; School of Chinese Medicine, Hong Kong Baptist University

## Abstract

Time-course single-cell RNA sequencing (scRNA-seq) data have been widely applied to reconstruct the cell-type-specific gene regulatory networks by exploring the dynamic changes of gene expression between transcription factors (TFs) and their target genes. The existing algorithms were commonly designed to analyze bulk gene expression data and could not deal with the dropouts and cell heterogeneity in scRNA-seq data. In this paper, we developed dynDeepDRIM that represents gene pair joint expression as images and considers the neighborhood context to eliminate the transitive interactions. dynDeepDRIM integrated the primary image, neighbor images with time-course into a four-dimensional tensor and trained a convolutional neural network to predict the direct regulatory interactions between TFs and genes. We evaluated the performance of dynDeepDRIM on five time-course gene expression datasets. dynDeepDRIM outperformed the state-of-the-art methods for predicting TF-gene direct interactions and gene functions. We also observed gene functions could be better performed if more neighbor images were involved.

## I. Introduction

Exploring gene expression data is a common approach to reconstruct the gene regulatory networks (GRNs), which represent the physical bindings between the transcription factors and their target genes. In the past decades, many algorithms have been proposed to infer such TF-gene interactions on bulk gene expression data [1]–[4], which ignores the cell-type-specific gene expression due to the assumption of homogeneity among cells. Single cell RNA sequencing (scRNA-seq) is an emerging technology that captures the gene expression profile for each cell rather than averaging them from bulk cells. In consideration of the cell dynamics over time, time-course scRNA-seq gene expression data is intrinsically much more informative and impressive than the static scRNA-seq data, particularly for the inference of putative regulatory signals. However, most existing methods for GRN reconstruction were designed for static scRNA-seq data [5], [7], [8] or required pseudotime ordered cells [9], [23]. The classical statistical measures, such as Pearson correlation (PCC) [19] and Mutual information (MI) [20], can be applied to time-course scRNA-seq data by simply concatenating the data from each time point, but they cannot deal with dropout issue and ignore the inner temporal relationship between TFs and genes. In a more recent study, TDL [6] encoded the expression profile of a gene pair into an image and collected the images over time as a three-dimensional (3D) tensor. The interaction of this gene pair was further predicted by the two models: “TDL-3D CNN” and “TDL-LSTM”. Based on the TF-gene interactions, TDL was also proved its ability in predicting the functions relating cell cycle, rhythm, immune and proliferation genes. Because TDL only considers the target TF-gene pairs and neglects the neighboring context, it cannot rule out the false positives due to the transitive interactions (Fig. 2A), which has been proved by our previous paper [8].

**Fig. 1.**
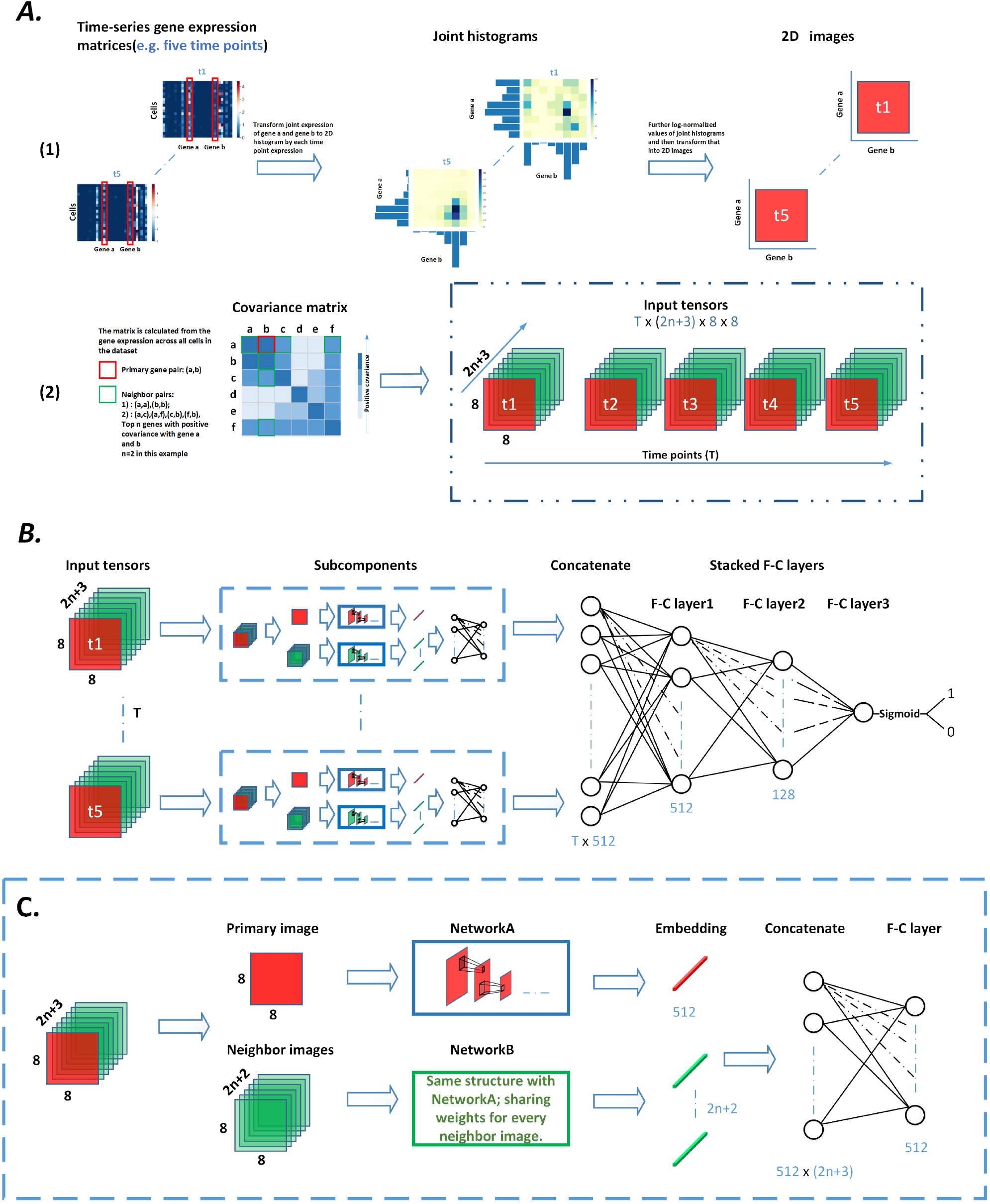
The overview and network structure of dynDeepDRIM. (A) The input of dynDeepDRIM as a four-dimensional (4D) tensor. (1) Representation of the joint gene expression of gene pair (*a, b*) as a primary image using joint histogram; (2) For each time point, the 2*n* + 2 neighbor images(green color) are generated from the joint gene expression of neighbor gene pairs (*a, a*), (*b, b*), (*a, c*), (*a, f*), (*c, b*), and (*f, b*). Next, the primary image and the neighbor images are concatenated as a three-dimensional (3D) images tensor as the representation of gene pair (*a, b*) in a time point. For *T* time points, the 3D tensors of each time point are stacked along time-axis into a 4D tensor as the input of dynDeepDRIM. (B) The structure of dynDeepDRIM. It consists of *T* subcomponents followed by three fully connected (F-C) layers to produce the prediction values using Sigmoid function for binary classification. (C) The network architecture of subcomponent. It includes NetworkA and NetworkB, which are used to process primary and neighbor images, respectively.

**Fig. 2.**
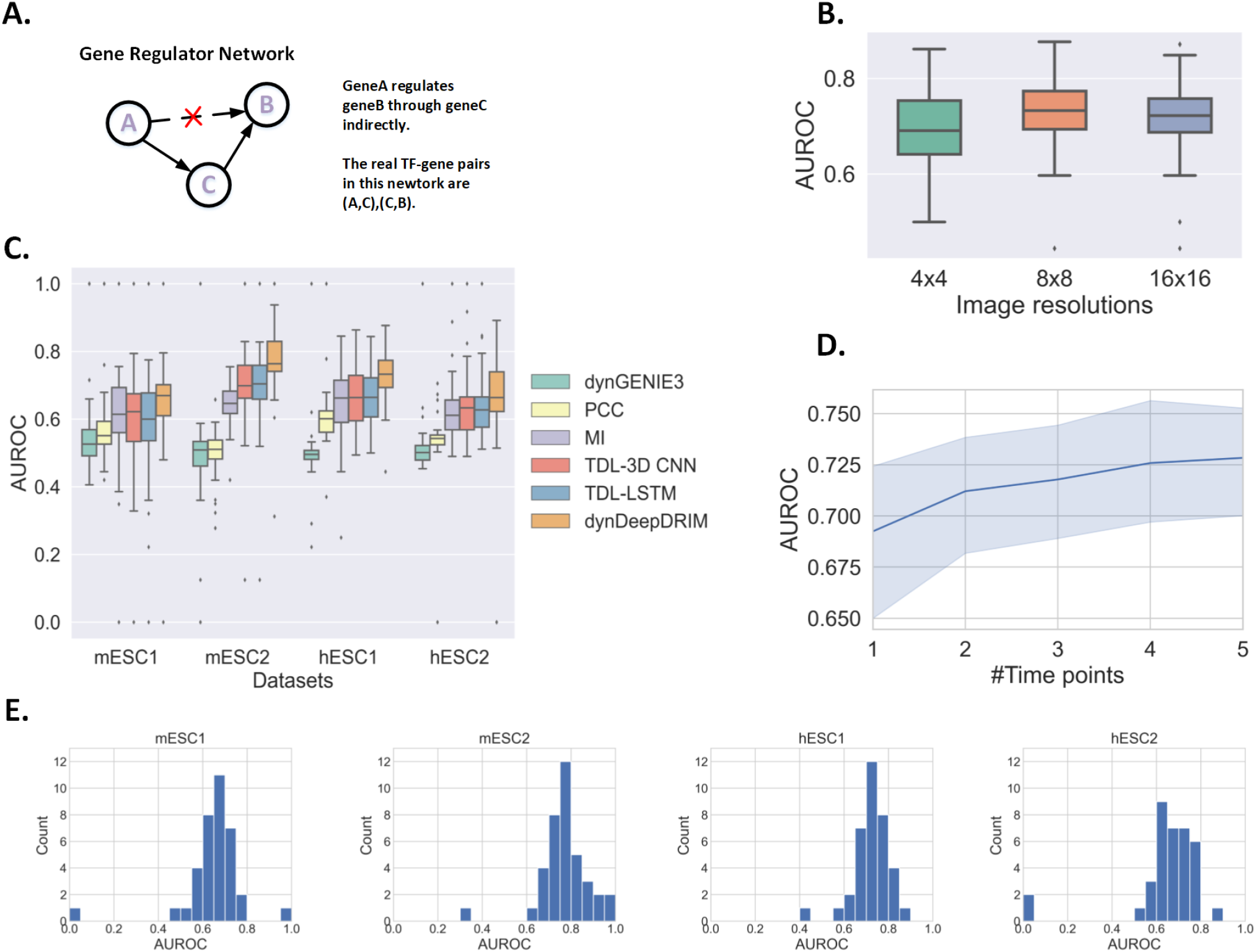
The predictive performance of dynDeepDRIM in TF-gene interactions. (A) An example of transitive interaction. (B) The influence of different image resolutions in hESC1 dataset with *T* = 5, *n* = 10. (C) The performance of the six algorithms on TF-gene regulatory interaction prediction. (D) The curve of the median AUROC by involving different time points in hESC1 dataset. The gray area represents the standard errors. (E) The distribution of AUROC for each TF calculated by dynDeepDRIM in the four datasets.

In this paper, we proposed dynDeepDRIM to reconstruct GRNs on time-course scRNA-seq data using high-dimensional convolutional neural network (CNN). Other than considering only the image of the target TF-gene pair (primary image) as an input, dynDeepDRIM also involves the images from the gene pairs which share one gene with the target pair (neighbor images, Fig. 1A(2)) in the model to capture neighboring context. For each time point, dynDeepDRIM constructs a subcomponent involving two networks (NetworkA and NetworkB) to process the primary image and neighbor images, respectively (Fig. 1C). A fully connected network collects the outputs from the subcomponents over time points to infer the direct regulatory interaction between TF-genes pairs. We tested dynDeepDRIM on four time-course scRNA-seq datasets and found it outperformed the other state-of-the-art algorithms. We also applied dynDeepDRIM to predict gene functions on mouse brain cortex dataset and found the neighbor images could not only remove transitive interactions from the predicted GRNs but also significantly improve the gene function annotations.

## II. Methods

The time-courses gene expression profiles are represented as a series of matrices 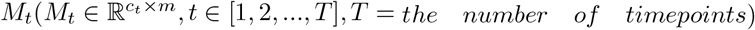, where *c*_*t*_ is the number of cells (rows) of *t*-th time point and *m* is the number of genes (columns). In data preprocessing, we normalized the raw read counts of gene expression into Reads Per Kilobase of transcript per Million mapped reads (RPKM). As shown in the leftmost panel of Fig. 1A(1), the heatmaps give an intuitive example of time-course gene expression profiles after normalization.

### A. Represent the joint expression of gene pairs as images

dynDeepDRIM represents the joint histograms of the expression for each gene pair as an image. It adds a pseudo-count (10^−2^) to all the entries in *M*_*t*_ to alleviate the influence of dropouts and applies log-normalization to *M*_*t*_ to avoid the extreme expression values. Denote 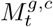 as the expression of gene *g* in cell *c* at *t*-th time point. The log-normalization computes using the following function:
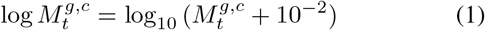

We assume the processed expression values of gene *a* and gene *b* are 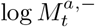 and 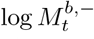 (′− ′ represents all of the cells in *M*_*t*_) and split the values in 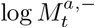 and 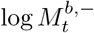 by equal-width 8 bins (by default) to generate their histograms as 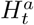 and 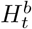. Their joint histogram 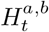 (8 by 8, shown in the middle panel of Fig. 1A(1)) is further log-normalized to avoid extreme values:
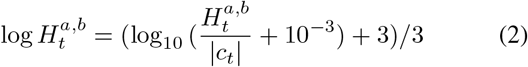

|*c*_*t*_| represents the number of cells in time point *t*, which is used here as a scaling factor to reduce the impact of different cell numbers over time points. An image with 8 × 8 pixels (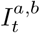, shown in the rightmost panel of Fig. 1A(1)) is generated to represent the joint expression of gene pair (*a, b*).

### B. Represent primary and neighbor images as a tensor

Assuming the target gene pairs are (*a, b*) and their expression profiles will be represented as the primary image at time point *t*. Similarly, we represent the neighborhood context of gene pair (*a, b*) as neighbor images, which encompass two parts: 1) self-images 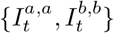; 2) 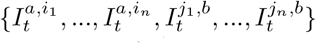, where the gene sets (*i*_1_, *i*_2_, …, *i*_*n*_) and (*j*_1_, *j*_2_, …, *j*_*n*_) are the top *n* genes with strong positive covariance to gene *a* and gene *b*, respectively. Thus, 2*n* + 2 neighbor images are generated for a gene pair (*a, b*) (Fig. 1A(2)). We stack the primary and the neighbor images into a 3D tensor ((2*n*+3) × 8 × 8) as the representation of gene pair (*a, b*) at time point *t*. dynDeepDRIM will collect the 3D tensors over time points and aggregate them as a 4D tensor as its input.

### C. Model structure and training strategy

As shown in Fig. 1B, the network structure of dynDeepDRIM includes *T* subcomponents to process the 3D tensors from different time points. Each subcomponent consists of two similar CNN networks (denoted as NetworkA and NetworkB) based on VGGnet [25] with non-linear activation function ReLU. NetworkA is used to process the primary image, embedding it into a vector of size 512 (by default, it is determined by the number of nodes in the fully connected layer), while NetworkB is a siamese-like network for embedding 2*n* + 2 neighbor images. The embeddings (512 × (2*n*+3)) of the 2*n*+3 images are transformed into another condensed embedding with 512 dimensions used to integrate with the results for the other time points by the three fully connected layers (Fig. 1B and C). The Sigmoid function is used to generate the final prediction score between 0 and 1 for binary classification (Fig. 1B). dynDeepDRIM is trained by mini-batched stochastic gradient descent with batch-size 32, and validation set is randomly selected 20% from training set for model selection and early stopping.

### D. Performance evaluation for GRN reconstruction

We adopted three-fold cross-validation to assess the performance of three supervised models (TDL-3D CNN, TDL-LSTM and dynDeepDRIM) in the four time-course scRNA-seq data for GRN reconstruction (**Results**). We kept balanced positive and negative pairs for each TF and divided all the TFs into three partitions. We carefully adjusted the assignment of TFs to make sure the numbers of TF-gene pairs are close among partitions. For three-fold cross-validation, the model was trained using the TF-gene pairs from two partitions, and tested on the one from the remaining partition.

### E. Performance evaluation for gene function annotation

We extracted the intersection between 1. the genes with specific functions; and 2. the ones from time-course scRNA-seq data as positive cases (*K*) and randomly selected the remaining genes from scRNA-seq data as negative cases (*U*, |*K*| = |*U*|). We selected 2*/*3 of the genes in *K* and *U* as training set (denoted as *K*_*train*_ and *U*_*train*_), and the remaining 1*/*3 was test set (denoted as *K*_*test*_ and *U*_*test*_). The labels of gene pairs were generated based on the following rules:

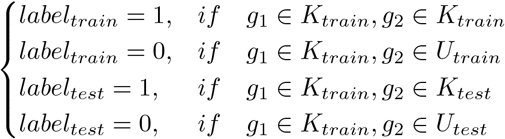

where *g*_1_ and *g*_2_ represent a genes pair. By considering the gene pairs, the size of training set is |*K*_*train*_| × |*K*_*train*_| + |*K*_*train*_| × |*U*_*train*_|, and it is |*K*_*train*_| × |*K*_*test*_| + |*K*_*train*_| × |*U*_*test*_| for the testing set.

## III. Results

### A. Time-course scRNA-seq datasets

In this study, we downloaded four time-course scRNA-seq datasets from Gene Expression Omnibus [10] and European Molecular Biology Laboratory-European Bioinformatics Institute(EMBL-EBI), two from mouse (mESC1 and mESC2) and two from human (hESC1 and hESC2) embryonic stem cells [11]–[14]. Besides the above mentioned four scRNA-seq datasets, the mouse brain time-course scRNA-seq dataset [15] was used to evaluate the performance of dynDeepDRIM for gene function annotation (Table. I).

**TABLE I.** Summary for time-course scRNA-seq datasets for experiment

| Datasets | Number of Cells | Number of Time points | Accession number |
| --- | --- | --- | --- |
| mESC1 | 3,456 | 9 | GSE79578 |
| mESC2 | 2,717 | 4 | GSE65525 |
| hESC1 | 1,529 | 5 | E-MTAB-3929 |
| hESC2 | 758 | 6 | GSE75748 |
| Mouse brain cortex | 21566 | 3 | GSE104158 |

### B. Benchmark for regulatory interactions and gene functional similarities

#### 1) The benchmark for GRN reconstruction

We collected 38 TFs for mESC2 and 36 TFs for the other three datasets. Their potential targets were inferred from the ChIP-seq data (*P*-value < 10^−400^ with peak signals) in the gene transcription regulation database(GTRD) [21] as benchmarks to define positive and negative pairs. The positive pairs were defined as the TF had one or more significant peak signals in the promoter region of gene *b*. The gene promoter regions were defined as 10Kb upstream or 1Kb downstream of the gene transcription start site [6], [24].

#### 2) The benchmark for gene annotation

We downloaded the four gene sets from GSEA-MsigDB [16]–[18], they were cell cycle (614 genes, GO:0044770), immune (332 genes, GO:0002376), rhythm (207 genes, WP3594), and proliferation (138 genes, GO:0061351). The positive and negative gene pairs in training and test sets were generated based on the approach described in section II-E.

### C. Determine image resolution

The image resolution for each gene pair is a hyperparameter and could be fine-tuned by dynDeepDRIM. We compared the performance of dynDeepDRIM with image resolutions of 4 × 4, 8 × 8, and 16 × 16 in hESC1 dataset and found their performance was not changed dramatically (Fig. 2B). We chose the best resolution of 8 × 8 in the experiment.

### D. Performance on the four time-course scRNA-seq datasets for TF-gene interactions

We evaluated the performance of dynDeepDRIM to predict TF-gene direct regulatory interactions using four time-course scRNA-seq datasets and compared it with dynGENIE3 [22], PCC [19], MI [20], TDL-3D CNN and TDL-LSTM [6]. Because PCC and MI were not designed for time-course gene expression data, we merged the cells from different time points in the same matrix. dynGENIE3 was designed for bulk time-course gene expression data, so we averaged the gene expression values among the cells for each time point. As the results shown in Fig. 2C, we observed dynDeepDRIM outperformed the other five algorithms and the supervised methods(dynDeepDRIM, TDL-3DCNN and TDL-LSTM) were better than the unsupervised ones (dynGENIE3, PCC and MI) in most of the datasets. We noticed there were some TFs with outlier AUROC values, that because these TFs have extremely few target genes in the benchmark data (e.g. *pou*3*f*1 and *smarcc*1 only have one target gene in mESC1 and mESC2). We also showed the histograms of TF-specific AUROC in the four datasets (Fig. 2E).

### E. Influence of the number of time points

We generated five datasets with the subsets of consecutive time points ({*t*1}, {*t*1, *t*2}, …, {*t*1, *t*2, *t*3, *t*4, *t*5}) of hESC1 to evaluate the influence of the number of time points by dynDeepDRIM. As shown in Fig. 2D, we found the performance of dynDeepDRIM consistently increased by involving more time points.

### F. Gene function prediction

dynDeepDRIM can also be used to annotate gene functions by assuming the genes shared the same biological functions if they have direct interactions. We used the time-course scRNA-seq data from mouse brain (Table. I) and downloaded four functional annotated gene sets from GSEA-MsigDB [16]. We extracted their shared genes for the corresponding gene functions, cell cycle (568 genes), immune (269 genes), rhythm (187 genes) and proliferation (127 genes). We observed dynDeepDRIM significantly outperforms the two models in TDL for all the datasets(Fig. 3A; Average AUROC=0.91,0.66,0.66 for dynDeepDRIM, TDL-3D CNN and TDL-LSTM, respectively). Interestingly, we explored the AUROCs were increased by incorporating more neighbor images and the trends were consistent for all the four functional annotations (Fig. 3B, C and D).

**Fig. 3.**
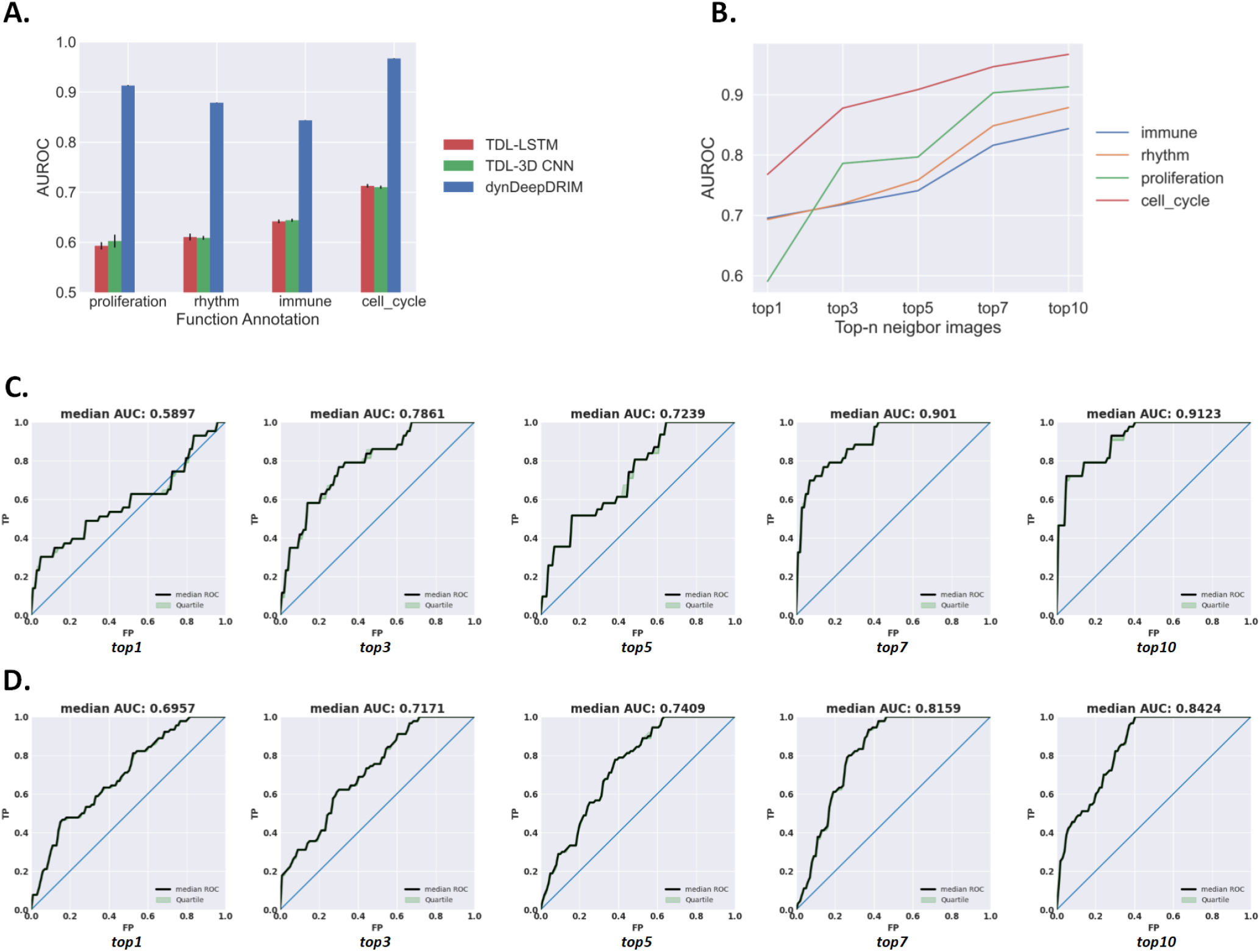
The predictive performance of dynDeepDRIM in gene functional annotation. (A) Comparison of TDL-LSTM, TDL-3D CNN and dynDeepDRIM on gene functional annotation. (B) The influence of involving different neighbor images. (C) The ROC curves of dynDeepDRIM for proliferation annotation with different neighbor images. (D) The ROC curves of dynDeepDRIM for immune annotation with different neighbor images.

## IV. Discussion and conclusion

Time-course scRNA-seq is a powerful tool to capture the dynamic changes of gene expression over time points, which is important to interpret disease progression or organ development. Rather than the impact of single gene, GRNs defines the synergic effects between TFs and genes to implement the specific biological functions. Exploring such TF-gene interactions is expensive and with low-throughput using wetlab experiments, preventing us to understand the whole network system. Bulk gene expression data has been used to reconstruct GRNs using computational methods for a long time due to the fact that TFs and their target genes commonly co-express in transcriptional level. The emerging scRNA-seq technology challenges these algorithms by introducing cell-cell heterogeneity and dropouts. Although the algorithms, such as CNNC [7], SCODE [23], has been developed to predict TF-gene interactions on static scRNA-seq data, but the time-course scRNA-seq data was seldomly considered. TDL was recently introduced to predict TF-gene interactions in time-course scRNA-seq data, but such strategy would introduce considerable false positives due to the transitive interactions [8].

In this study, we proposed dynDeepDRIM, a deep neural network, to predict TF-gene interactions on time-course scRNA-seq data. Rather than consider only the target TF-gene pairs, dynDeepDRIM also considers the neighbor context to distinguish the transitive interactions. We observed it significantly outperformed the other methods and noticed the number of time points could positively influence the predictive performance for the particular datasets. More time points would result in involving more cells in the prediction model, which could influence the optimal image resolution. It does not always true that high resolution leads to better predictions, because it is sensitive to noise and make the expression histogram unstable. Because the GRN is commonly cell-type-specific, it should be also noticed to select the matched ChIP-seq data as training labels. Besides GRN reconstruction, dynDeepDRIM could also help in gene function prediction by assuming the genes with direct interactions sharing the same biological functions. We identified involving the neighbor images can also help this task, because the two genes with the same function commonly share the same neighbor genes. For example, *mia*3 and *il*6*st* are both with “immune” function and they share 7 neighbor genes (*n* = 10). This observation motivates us to consider the neighbor images for gene function annotations.

## Competing interests

The authors declare that they have no competing interests.

## Additional Files

The codes of dynDeepDRIM are available at: https://github.com/yuxu-1/dynDeepDRIM.

## Authors’ contributions

LZ conceived the study; YX, JXC designed dynDeepDRIM; YX implemented the algorithm and analyzed the results. YX, JXC conducted the experiments. YX, LZ wrote the article. APL and WC reviewed the paper. All authors read and approved the final manuscript.

## Funding

This research is partially supported by Hong Kong Research Grant Council Early Career Scheme (HKBU 22201419), HKBU Start-up Grant Tier 2(RC-SGT2/19-20/SCI/007), HKBU IRCMS (No. IRCMS/19-20/D02) and Guangdong Basic and Applied Basic Research Foundation (No. 2019A1515011046 and No. 2021A1515012226). SZVUP Special Fund Project (2021Szvup135).

## Acknowledgment

The authors would like to thank Research Grants Council of Hong Kong, Hong Kong Baptist University and HKBU Research Committee for their kind support of this project.

## Notes

### Competing Interest Statement

The authors have declared no competing interest.

## References

[1] Haury A C, Mordelet F, Vera-Licona P, et al., TIGRESS: trustful inference of gene regulation using stability selection[J]. BMC systems biology, 2012, 6(1): 1–17.

[2] Faith J J, Hayete B, Thaden J T, et al., Large-scale mapping and validation of Escherichia coli transcriptional regulation from a compendium of expression profiles[J]. PLoS biology, 2007, 5(1): e8.

[3] Kuüffner R, Petri T, Tavakkolkhah P, et al., Inferring gene regulatory networks by ANOVA[J]. Bioinformatics, 2012, 28(10): 1376–1382.

[4] Huynh-Thu V A, Irrthum A, Wehenkel L, et al., Inferring regulatory networks from expression data using tree-based methods[J]. PloS one, 2010, 5(9): e12776.

[5] Pratapa A, Jalihal A P, Law J N, et al., Benchmarking algorithms for gene regulatory network inference from single-cell transcriptomic data[J]. Nature methods, 2020, 17(2): 147–154.

[6] Ye Yuan, Ziv Bar-Joseph, Deep learning of gene relationships from single cell time-course expression data. Briefings in Bioinformatics, 2021. https://doi.org/10.1093/bib/bbab142

[7] Yuan Y, Bar-Joseph Z, Deep learning for inferring gene relationships from single-cell expression data[J]. Proceedings of the National Academy of Sciences, 2019, 116(52): 27151–27158.

[8] Chen, Jiaxing, et al., DeepDRIM: a deep neural network to reconstruct cell-type-specific gene regulatory network using single-cell RNA-seq data.. bioRxiv (2021).

[9] Zhang Y, Chang X, Liu X, Inference of gene regulatory networks using pseudo-time series data[J]. Bioinformatics, 2021.

[10] Edgar R, Domrachev M, Lash A E, Gene Expression Omnibus: NCBI gene expression and hybridization array data repository[J]. Nucleic acids research, 2002, 30(1): 207–210.

[11] Semrau S, Goldmann JE, Soumillon M, Mikkelsen TS et al., Dynamics of lineage commitment revealed by single-cell transcriptomics of differentiating embryonic stem cells. Nat Commun 2017 Oct 23;8(1):1096. PMID: 29061959

[12] Klein AM, Mazutis L, Akartuna I, Tallapragada N et al., Droplet barcoding for single-cell transcriptomics applied to embryonic stem cells. Cell 2015 May 21;161(5):1187-1201. PMID: 26000487

[13] Petropoulos S, Edsgard D, Reinius B, et al., Single-cell RNASeq reveals lineage and X chromosome dynamics in human preimplantation embryos. Cell 2016;165:1012–26

[14] Chu LF, Leng N, Zhang J, Hou Z et al., Single-cell RNA-seq reveals novel regulators of human embryonic stem cell differentiation to definitive endoderm. Genome Biol 2016 Aug 17;17(1):173. PMID: 27534536

[15] Mayer C, Hafemeister C, Bandler RC, Machold R et al., Developmental diversification of cortical inhibitory interneurons. Nature 2018 Mar 22;555(7697):457-462. PMID: 29513653

[16] Subramanian A, Tamayo P, Mootha V K, et al., Gene set enrichment analysis: a knowledge-based approach for interpreting genome-wide expression profiles[J]. Proceedings of the National Academy of Sciences, 2005, 102(43): 15545–15550.

[17] Liberzon A, Birger C, Thorvaldsdóttir H, et al., The molecular signatures database hallmark gene set collection[J]. Cell systems, 2015, 1(6): 417–425.

[18] Liberzon A, Subramanian A, Pinchback R, et al., Molecular signatures database (MSigDB) 3.0[J]. Bioinformatics, 2011, 27(12): 1739–1740.

[19] Salleh F H M, Arif S M, Zainudin S, et al., Reconstructing gene regulatory networks from knock-out data using Gaussian Noise Model and Pearson Correlation Coefficient[J]. Computational biology and chemistry, 2015, 59: 3–14.

[20] Song L, Langfelder P, Horvath S, Comparison of co-expression measures: mutual information, correlation, and model based indices[J]. BMC bioinformatics, 2012, 13(1): 1–21.

[21] Yevshin I, Sharipov R, Valeev T, et al., GTRD: a database of transcription factor binding sites identified by ChIP-seq experiments. Nucleic Acids Res 2017;45:D61–7.

[22] Huynh-Thu, V., Geurts, P, dynGENIE3: dynamical GENIE3 for the inference of gene networks from time series expression data. Sci Rep 8, 3384 (2018) https://doi.org/10.1038/s41598-018-21715-0.

[23] Matsumoto H, Kiryu H, Furusawa C, et al., P, SCODE: an efficient regulatory network inference algorithm from single-cell RNA-Seq during differentiation[J]. Bioinformatics, 2017, 33(15): 2314–2321.

[24] Ernst J, Plasterer H L, Simon I, et al., Integrating multiple evidence sources to predict transcription factor binding in the human genome[J]. Genome research, 2010, 20(4): 526–536.

[25] Simonyan K, Zisserman A, Very deep convolutional networks for largescale image recognition[J]. arXiv preprint 1409.1556, 2014.

